# Cytoplasmic tail truncation of SARS-CoV-2 Spike protein enhances titer of pseudotyped vectors but masks the effect of the D614G mutation

**DOI:** 10.1101/2021.06.21.449352

**Authors:** Hsu-Yu Chen, Chun Huang, Lu Tian, Xiaoli Huang, Chennan Zhang, George N. Llewellyn, Geoffrey L. Rogers, Kevin Andresen, Maurice R.G. O’Gorman, Ya-Wen Chen, Paula M. Cannon

## Abstract

The high pathogenicity of SARS-CoV-2 requires it to be handled under biosafety level 3 conditions. Consequently, Spike protein pseudotyped vectors are a useful tool to study viral entry and its inhibition, with retroviral, lentiviral (LV) and vesicular stomatitis virus (VSV) vectors the most commonly used systems. Methods to increase the titer of such vectors commonly include concentration by ultracentrifugation and truncation of the Spike protein cytoplasmic tail. However, limited studies have examined whether such a modification also impacts the protein’s function. Here, we optimized concentration methods for SARS-CoV-2 Spike pseudotyped VSV vectors, finding that tangential flow filtration produced vectors with more consistent titers than ultracentrifugation. We also examined the impact of Spike tail truncation on transduction of various cell types and sensitivity to convalescent serum neutralization. We found that tail truncation increased Spike incorporation into both LV and VSV vectors and resulted in enhanced titers, but had no impact on sensitivity to convalescent serum inhibition. In addition, we analyzed the effect of the D614G mutation, which became a dominant SARS-CoV-2 variant early in the pandemic. Our studies revealed that, similar to the tail truncation, D614G independently increases Spike incorporation and vector titers, but that this effect is masked by also including the cytoplasmic tail truncation. Therefore, the use of full-length Spike protein, combined with tangential flow filtration, is recommended as a method to generate high titer pseudotyped vectors that retain native Spike protein functions.

**IMPORTANCE:** Pseudotyped viral vectors are useful tools to study the properties of viral fusion proteins, especially those from highly pathogenic viruses. The Spike protein of SARS-CoV-2 has been investigated using pseudotyped lentiviral and VSV vector systems, where truncation of its cytoplasmic tail is commonly used to enhance Spike incorporation into vectors and to increase the titers of the resulting vectors. However, our studies have shown that such effects can also mask the phenotype of the D614G mutation in the ectodomain of the protein, which was a dominant variant early in the COVID-19 pandemic. To better ensure the authenticity of Spike protein phenotypes when using pseudotyped vectors, we therefore recommend using full-length Spike proteins, combined with tangential flow filtration methods of concentration, if higher titer vectors are required.

## INTRODUCTION

Coronavirus disease 2019 (COVID-19) is caused by severe acute respiratory syndrome coronavirus 2 (SARS-CoV-2) and was first reported in Wuhan, China, in December 2019 (1). The disease rapidly spread worldwide, causing over 150 million confirmed cases and more than 3 million reported deaths by May 2021 (2). The accompanying worldwide research effort has resulted in a large number of vaccine candidates, and both national and international clinical trials to assess novel and repurposed drug regimens (3). In the United States, the SARS-CoV-2 Spike glycoprotein has been a primary target of such efforts. Spike is a major viral antigen that induces protective immune responses in COVID-19 (4–6) patients and mediates cell entry by binding to angiotensin-converting enzyme 2 (ACE2) (1, 7, 8) or other receptors (9, 10). ACE2 is expressed in the human respiratory system (11), especially on type II pneumocytes (12), which are the main target cell for SARS-CoV-2 infection. Expression of ACE2 in other organs also allows infection outside the lung (11).

Due to the high pathogenicity of SARS-CoV-2, biosafety level 3 (BSL3) labs are required for studies that involve replication-competent virus. Therefore, investigators often use Spike protein pseudovirus vector systems, based on replication-incompetent vector particles and attenuated or conditional viruses. Identification of an optimal pseudovirus system for any particular viral entry glycoprotein typically involves comparing the most commonly used systems: replication-incompetent lentiviral (LV) or retroviral (RV) vectors, or conditional vesicular stomatitis virus (VSV) viruses that are deleted for the VSV glycoprotein (G) (13, 14). SARS-CoV-2 Spike protein can pseudotype all three vector systems, which have been used to investigate viral entry (15–18), neutralization by monoclonal antibodies or convalescent plasma (4, 5, 15, 19–27), entry inhibitors (15, 23, 28, 29), and to characterize surging viral variants (30–44). In approximately one third of these studies, deletions of 18-21 amino acids from the cytoplasmic tail of Spike were used to enhance vector titers and thereby facilitate the study.

Cytoplasmic tail truncation of viral glycoproteins is a common strategy to enhance pseudovirus formation since this can remove steric interference that may occur between the heterologous viral glycoproteins and the vector matrix or capsid proteins (45–50). Also employed are cytoplasmic tail swaps, whereby the tail from the natural viral glycoprotein is used to create a chimeric glycoprotein with enhanced incorporation properties (46, 51). However, we and others have shown that tail modifications can also have functional consequences, for example, removing endocytosis signals that lead to increased cell surface levels and enhanced incorporation into vector particles (52, 53), alterations of the ectodomain conformation (52, 54), changes to fusogenicity (46, 54–56) and altered antigenic characteristics (57, 58).

SARS-CoV-2 Spike contains a putative ER retention signal (KLHYT) at its C-terminus, which is removed by the tail truncations of 13 amino acids (59) or 18-21 amino acids that are frequently employed (24, 60–63). Compared to the full-length Spike, such truncations were reported to generate ~10-20-fold higher titers of both LV vectors (59–61, 63) and VSV vectors (24, 60, 62). Truncated Spike also enhances RV vector titers, albeit with a smaller effect when compared side by side with LV and VSV vectors (60). Havranek *et al.* (62) and Yu et al. (59) investigated the mechanism for such an effect for VSV and LV pseudoviruses, and found that tail truncation enhanced both Spike incorporation into the viral particles and cell-cell fusion for Spike-expressing cells, but without altering cell surface expression levels. However, the impact of Spike tail truncations on any other ectodomain functions remains unclear.

In this report, we compared the practicality and functionality of using Spike pseudotyped vectors based on LV and VSV, and pseudotyped with either full-length or tail truncated proteins. We compared methods to prepare such vectors and identified tangential flow filtration as a facile method that is superior to ultracentrifugation and allows efficient production at a larger scale. An optimized system based on VSV vectors was used to assess the impact of the Spike mutation D614G (34), and to assess neutralizing activity in convalescent serum. Our studies determined that although Spike tail truncation boosts incorporation into vectors and enhances the titers achieved for unconcentrated supernatants, it also blunted the ability to observe differences caused by this specific Spike mutation. We therefore recommend that studies using Spike pseudotyped vectors retain the natural full-length cytoplasmic tail and use other strategies, such as concentration method and vector system choice, to achieve the required vector titers.

## RESULTS

### Spike cytoplasmic tail truncation facilitates vector incorporation and enhances titer

Cytoplasmic tail truncation of SARS-CoV-2 Spike has been reported to enhance the transduction efficiency of pseudotyped LV and VSV vectors (59–61, 63) with the effect suggested to be the result of enhanced incorporation and/or fusogenicity of Spike (59, 62). Using both the full-length Spike (S) and an 18 amino acid cytoplasmic tail truncation (SΔ18), we generated pseudotyped LV and VSV vectors carrying reporter GFP or luciferase genes, respectively. We compared the ability of the two Spike proteins to be incorporated into the vectors and to transduce HeLa cells expressing human ACE2 (HeLa-ACE2). Transduction by LV-GFP vectors was analyzed at 48 hours post-transduction, while VSV-Luc vectors were analyzed as early as 16-24 hours post-transduction.

Consistent with previous findings, we found that the cytoplasmic tail truncation increased vector transduction efficiency on HeLa-ACE2 cells, by approximately 4- and 30-fold for the LV-GFP and VSV-Luc vectors, respectively (Fig. 1A). We also observed a significant increase in incorporation for the truncated Spike protein in both vector systems (Fig. 1B), while having no impact on other viral particle proteins (Fig. 1B) or vector genome copy number (Fig. 1C). Together, these results suggest that cytoplasmic tail truncation increases Spike incorporation into both LV and VSV particles and this results in higher infectivity per particle. Since the VSV-Luc vectors have a faster read-out time, we chose this pseudovirus system for the rest of our studies.

**Figure 1.**
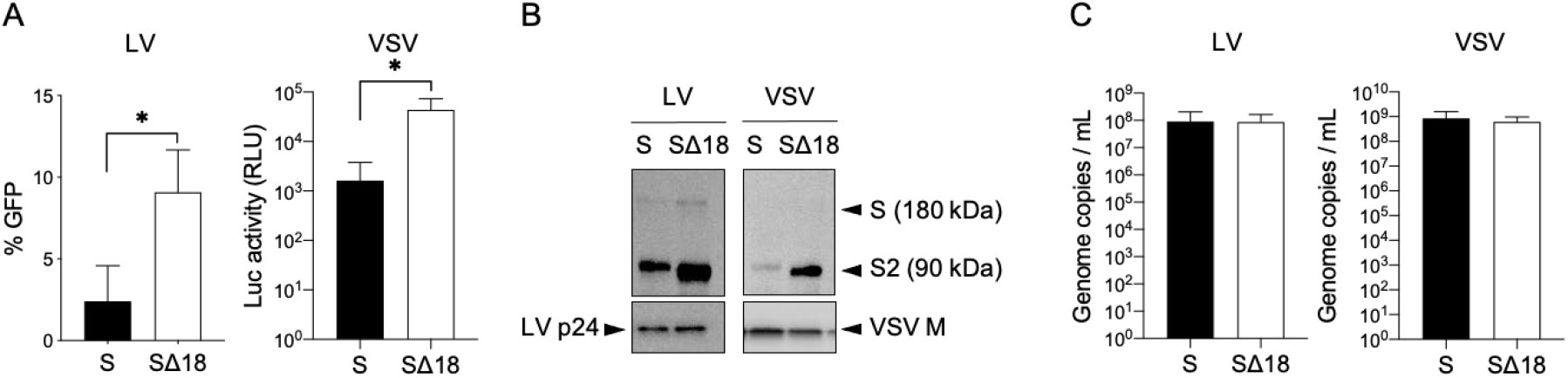
Impact of Spike protein cytoplasmic tail truncation on LV and VSV vectors. (A) Transduction of HeLa-ACE2 cells by equal volumes of unconcentrated vector supernatants of LV-GFP or VSV-Luc vectors, pseudotyped with full-length (S) or truncated (SΔ18) Spike proteins. Shown are mean and standard deviations from 3 independent vector stocks, *p<0.05, unpaired t-test, one-tail (B) Spike protein incorporation into vector particles, analyzed by Western blot using antibodies against the Spike S2 subunit and vector particle components p24 (LV) and M (VSV). Full length Spike (S) and S2 subunit are indicated. (C) Genomic copy number for indicated vectors. Shown are mean and standard deviations from 3 independent vector stocks.

### Susceptibility of different cell lines and lung organoids to Spike protein pseudovectors

Next, we tested the permissivity of different cell lines and a lung organoid model to SΔ18 pseudotyped VSV vectors. In agreement with previous finding, several ACE2-expressing cells were found to be susceptible to the vectors (1, 18), while ACE2 over-expression was required to support transduction of HeLa cells (Fig. 2A). We also evaluated an alternative transduction protocol with a shortened timeline, whereby trypsinized cells are incubated with the vectors simultaneously with seeding (64). Although this method shortened the overall process compared to a typical protocol that first seeds the cells in a tissue culture plate for 24 hours before incubation with vectors, it resulted in significantly lower transduction rates (Fig. 2B). Examination of cell surface ACE2 levels by flow cytometry revealed that newly trypsinized cells had 1.5-fold lower ACE2 levels compared to cells allowed to recover for 6 hours post-trypsinization (Fig. 2C), suggesting the reason for the lower titers.

**Figure 2.**
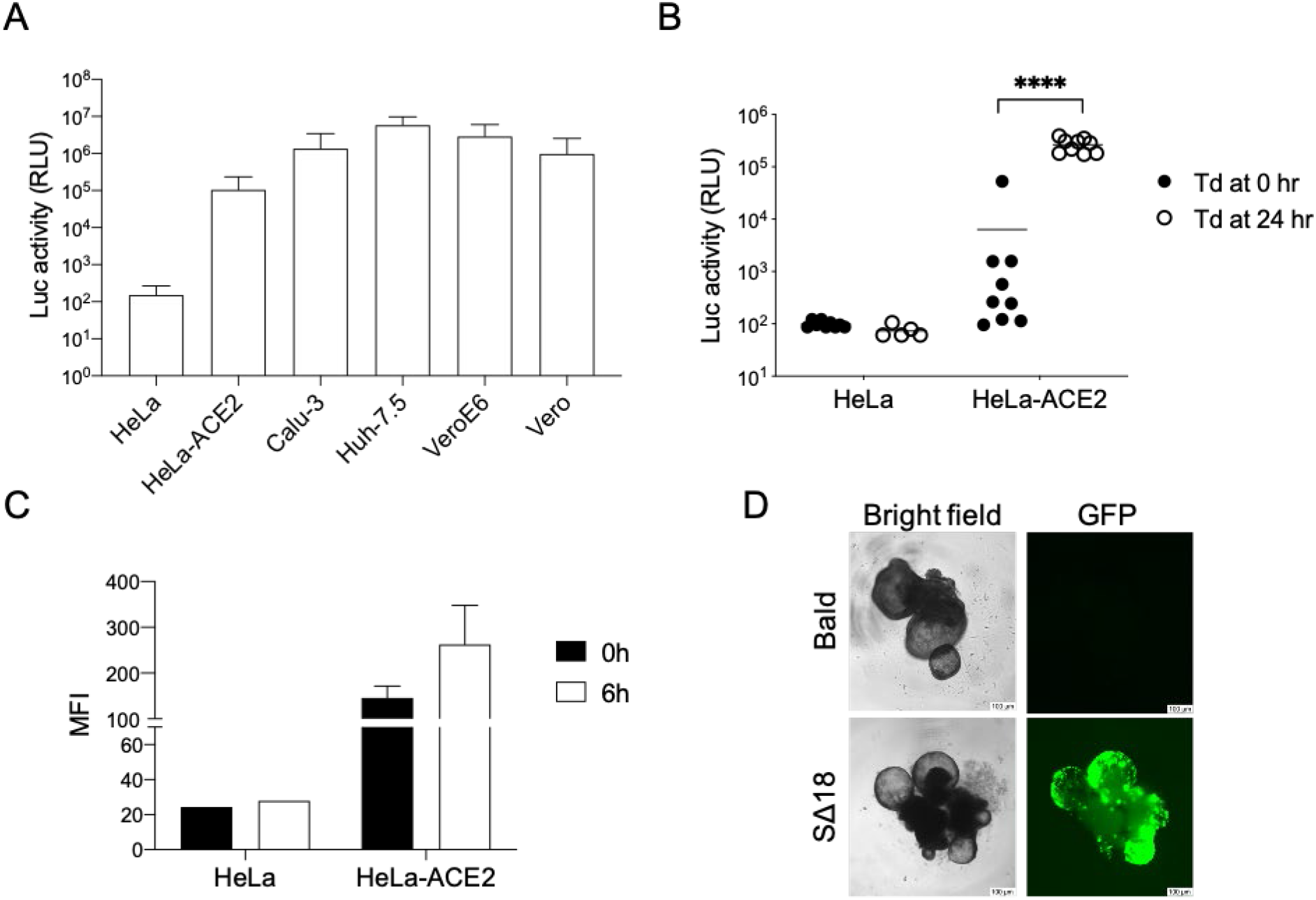
Transduction of cells by Spike VSV pseudovectors. (A) Indicated cell lines were transduced with equal amounts of SΔ18 VSV-Luc vectors and luciferase activity in cell lysates analyzed 24 hr later. Shown are mean and standard deviations from 3 independent vector stocks. (B) HeLa and HeLa-ACE2 cells were detached from culture flasks by trypsin, seeded into 96 well plates and transduced (Td) with equal amounts of SΔ18 VSV-Luc vectors. either immediately (0 hr) or 24 hours after seeding, and luciferase measured 24 hours later. Data from 9 different wells in a single experiment are shown. ****p<0.001, multiple T test. (C) ACE2 expression levels on cell surface measured by flow cytometry. Cells were stained with anti-ACE2 antibody at 0 or 6 hours after trypsinization. Means and standard deviations for MFI from two independent experiments are shown. (D) Lung bud organoids were transduced with equal amounts of VSV-GFP vectors pseudotyped with SΔ18 or control (bald) vectors with no glycoprotein. GFP expression was visualized 24 hours later.

Finally, we tested the susceptibility of a 3D lung bud organoid model to SΔ18 VSV pseudovectors carrying a GFP reporter. Compared to cell lines, lung organoids provide more physiologically relevant models of virus infection and have been used to identify candidate COVID-19 therapeutics (29). SΔ18 pseudotyped VSV-GFP vectors were able to efficiently transduce the cells, with GFP expression observed throughout the organoid by 24 hours (Fig. 2D).

### Tangential flow filtration facilitates scale-up of vector production and concentration

To identify an optimal method for concentration of Spike protein pseudovectors suitable for a research laboratory, we compared ultracentrifugation through a 20% w/v sucrose cushion with tangential flow filtration (TFF). Ultracentrifugation is limited by the capacity of a rotor, for example SW28 rotors have a maximum capacity of ~230 ml of vector supernatant per 2 hours run. In contrast, TFF can process much larger volume (65, 66) and a single TFF filter with 1000 cm^2^ surface area can process up to 3000 ml in 2 hours. Larger filter systems with capacities up to 15 L are also available. In addition, VSV-G pseudotyped LV vectors produced by TFF are reported to have a higher recovery rate than when prepared by ultracentrifugation (66).

To compare these approaches, 100 ml of SΔ18 pseudotyped VSV-Luc vector supernatants were subjected to either ultracentrifugation (Ultra) or TFF and concentrated into a 8ml final stock (12x, v/v). Vector genome copies in the unconcentrated and 12x concentrated vector stocks were measured by ddPCR, which revealed slightly better recovery rates following TFF (~70%) compared to ultracentrifugation (~60%) (Fig. 3A and 3B). At the same time, the transduction efficiencies of the three vector stocks were measured on HeLa-ACE2 cells, using serially diluted vectors (1:5 to 1:450 dilutions) (Fig. 3C). At the higher dilution points, both 12x Ultra and 12x TFF vector preparations produced about a 10-fold higher luciferase signal compared to the unconcentrated vectors. Interestingly, at the lower dilutions (1:5 and 1:15), the 12x Ultra vector stock showed no enhancement over unconcentrated vectors, while the 12x TFF vector stocks retained their 10-fold higher transduction rates. This observation is suggestive of the presence of an inhibitory factor that is concentrated during ultracentrifugation but was not retained following TFF.

**Figure 3.**
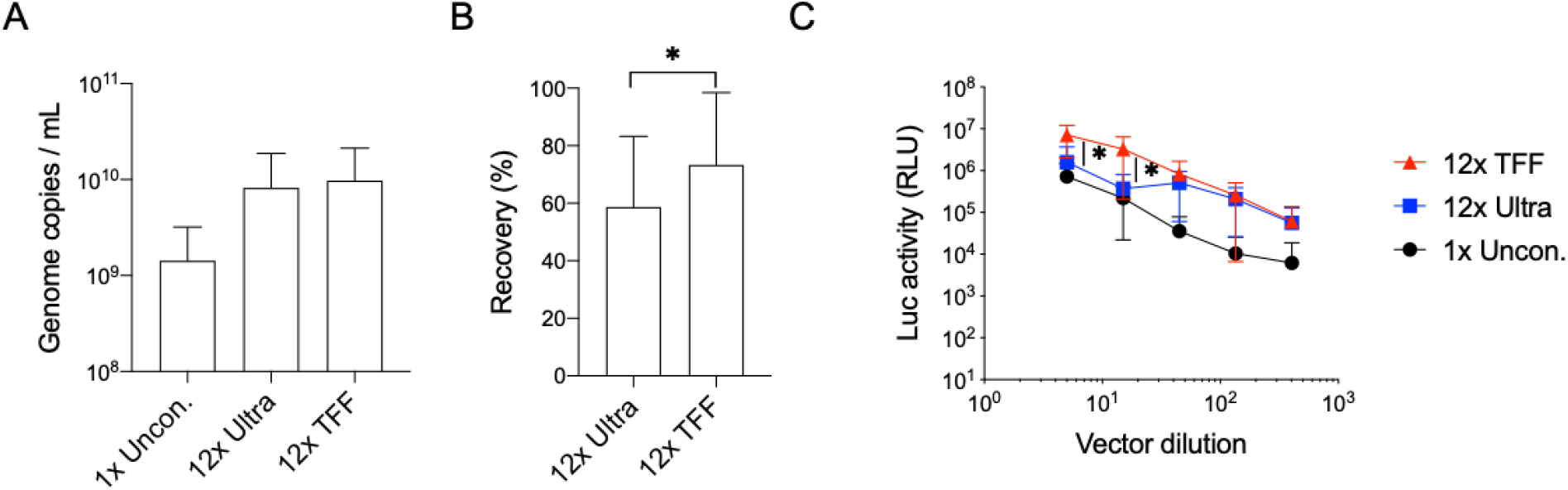
Concentration methods for Spike VSV pseudovectors. (A) Genome copy numbers of VSV-Luc vectors pseudotyped with SΔ18 Spike protein, from unconcentrated supernatants (1x Uncon.), or following 12x concentration (v/v) by either ultracentrifugation (Ultra) or tangential flow filtration (TFF). Shown are mean and standard deviations from 3 independent vector stocks. (B) Vector recovery, calculated by comparing genome copies in concentrated versus unconcentrated vector stocks. Shown are mean and standard deviation from 3 independent vector concentrations for each method, *p<0.05, one-tailed Paired T test. (C) Transduction of HeLa-ACE2 cells by serial dilutions (1:5 to 1:405) of indicated vectors. Shown are mean and standard deviation from three independent vector stocks. *p<0.01, two tail Paired T test, for comparison between 12x Ultra and 12x TFF at the same dilutions.

In summary, we found that TFF facilitates large-scale processing of vector stocks, with similar genomic copy number recovery rates as the more typical ultracentrifugation method. More importantly, TFF results in vector stocks that retain a more consistent titer throughout a broader range of different dilutions than those produced by ultracentrifugation.

### Cytoplasmic tail truncation alters Spike protein functional properties

We used the VSV pseudovirus system to examine the impact of the D614G mutation of Spike protein. This mutation was first detected in China and Germany in late January and became the dominant circulating variant of SARS-CoV-2 globally by April 2020 (34). The mutation has functional consequences for the virus, resulting in higher viral loads in the upper respiratory tract (34, 67). *In vitro* studies with SARS-CoV-2 revealed that the D614G mutation enhanced replication on human lung epithelial cells and primary airway tissue (41), and increased replication or transmissibility in human ACE2 transgenic mice and hamster models (41, 68, 69). Effects were also observed using Spike protein pseudoviruses, where the D614G mutation was reported to enhance Spike incorporation into vector particles, despite minimal or no effect on Spike expression in vector-producing cells (43, 70), and to increase transduction rates on various cell lines (30, 32, 34, 37, 43, 63, 71).

Since we had noted that the cytoplasmic tail truncation of Spike protein also increased incorporation rates and transduction efficiencies (Fig. 1), we next examined the impact of the D614G mutation in the context of both full-length and truncated Spike proteins. For the full-length Spike protein, we observed up to 18-fold higher transduction rates for the G614 variant on HeLa-ACE2 cells, with less striking effects on the other cell lines we tested. In contrast, transduction rates for the variants in the SΔ18 backbone showed minimal to no differences across the range of cell types tested (Fig. 4A). The discrepancy between the behavior of the full-length S and SΔ18 vectors occurred despite similar genome copy numbers (Fig. 4B), ruling out an effect on vector production. We also observed no differences in cell surface expression levels of Spike when comparing the different variants in vector-producing cells (Fig. 4C). Instead, in agreement with previous studies using full-length Spike pseudotyped RV and LV vectors, we found that the D614G mutation enhanced Spike incorporation, albeit with a much larger effect for the full-length Spike versus the truncated protein (~9-fold versus ~2-fold effect) (Fig. 4D, E). Together, these observations suggest that a primary effect of both the tail truncation and the D614G mutation is on Spike protein incorporation, which in turn leads to enhanced titers, and that an upper limit for these effects likely reduces the impact of the D614G mutation when combined with a tail truncation.

**Figure 4.**
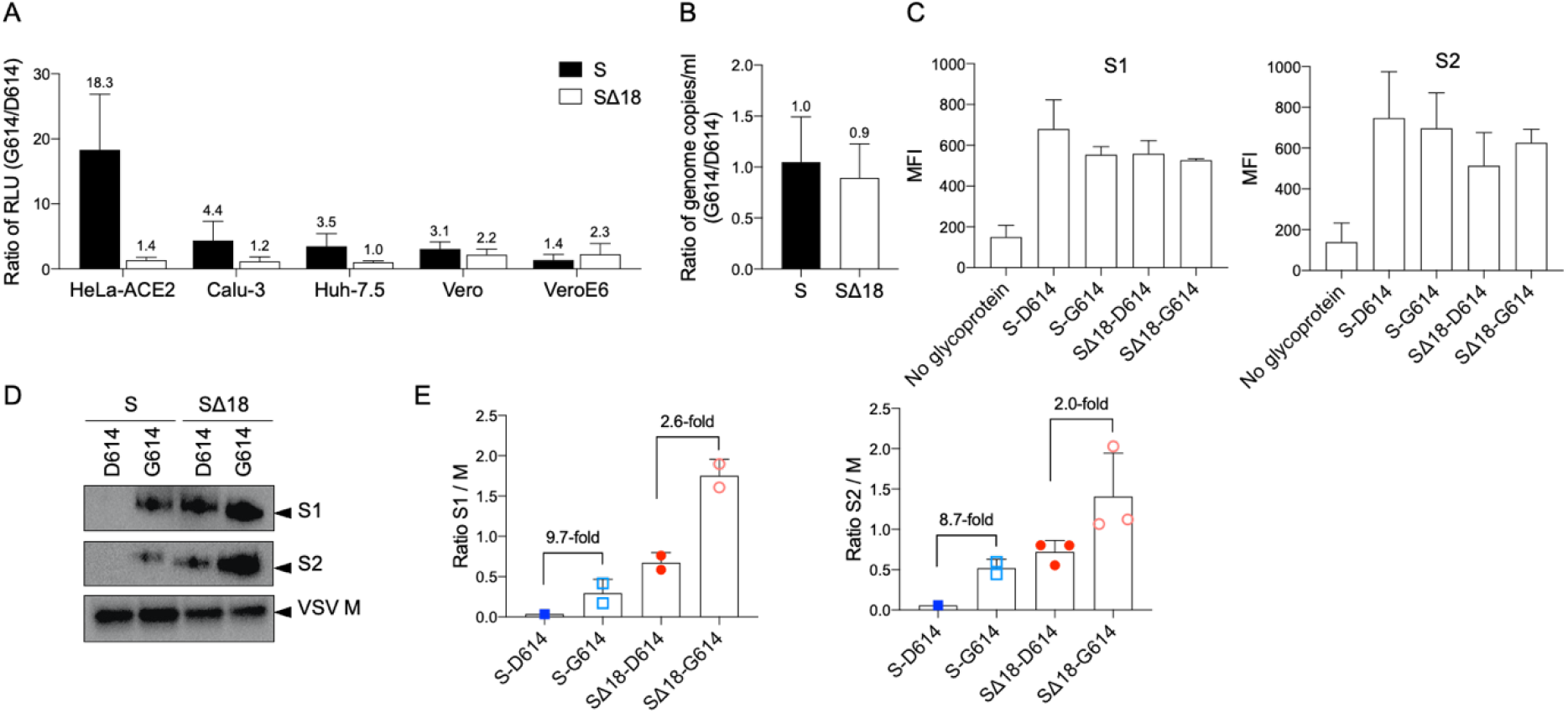
Impact of D614G mutation only observed with full-length Spike protein. (A) Indicated cell lines were transduced with G614 or D614 variants of VSV-Luc vectors, for both full-length and truncated Spike protein versions. The pairs of compared vectors were produced in the same way, and equal volumes applied. Luciferase activity was measured after 24 hours and ratios calculated. Means and standard deviations for 3 independent vector stocks are shown. (B) Ratio of genomic titers of G614 versus D614 vectors, for both full-length and truncated Spike proteins. Shown are mean and standard deviations from equal volumes of 3 independent vector stocks, produced in the same way for each pairwise comparison. (C) Cell surface expression levels of different Spike protein variants on 293T vector-producing cells, measured by flow cytometry using anti-S1 or anti-S2 antibodies at the time of vector harvest. Expression levels are reported as mean fluorescence intensity (MFI). Control 293T cells were from “bald” vector production, which were not transfected with any glycoprotein but still infected by the VSVΔG particles. Means and standard deviations from two independent experiments are shown. (D) Western blot showing incorporation of Spike proteins into VSV particles, from equal volumes of 100x concentrated vector supernatants, using antibodies against S1 or S2 subunits of Spike, or VSV M protein. (E) Comparison of Spike subunit incorporation into vectors, normalized to VSV M. Data from 2-3 independent vector stocks, indicated by individual dots. S1 and S2 subunits were only detected in one stock of S-D614 vectors.

### D614G mutation or cytoplasmic tail truncation does not alter Spike protein sensitivity to convalescent serum

Spike protein pseudovectors are a useful tool to measure antibody neutralizing activity in COVID-19 patient or convalescent sera (4, 5, 15, 19, 20, 23, 24, 26, 27, 61). We examined whether the D614G mutation or the cytoplasmic tail truncation altered sensitivity to neutralization by a panel of convalescent sera. VSV-Luc vectors displaying the four different Spike proteins were incubated with serially-diluted sera for 30 minutes before being applied to HeLa-ACE2 cells. After normalizing values to the luciferase signals obtained from cells transduced in the absence of sera, we observed that all four Spike proteins exhibited similar sensitivities to each serum (Fig 5). This suggests that neither the D614G mutation nor the cytoplasmic tail truncation alter the sensitivity of the Spike protein to neutralization.

**Figure 5.**
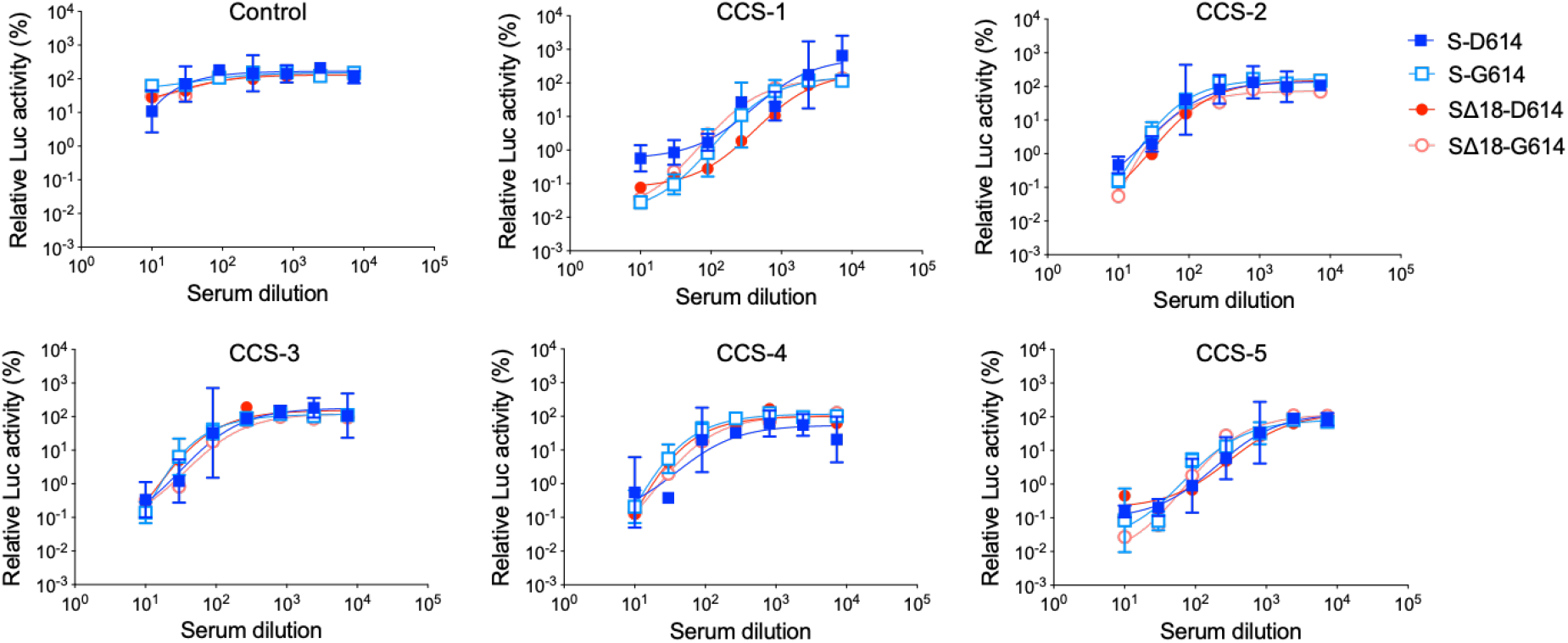
Sensitivity of different Spike proteins to convalescent serum neutralization. Indicated VSV-Luc pseudoviruses were incubated with serially-diluted sera (1:10 to 1:7290 fold) from control or convalescent COVID-19 patients (CCS) for 30 mins. before addition to HeLa-ACE2 cells. Luciferase activity was determined 24 hours later. All values were normalized to the luciferase signal from cells transduced with the same pseudovirus without serum. Means and standard deviations from 3 technical replicates of single vector stocks are shown.

## DISCUSSION

Pseudotyped vectors are a useful system to study the entry glycoproteins from highly pathogenic viruses such as SARS-CoV-2, as they remove the need for BSL-3 laboratory conditions. We confirmed that the SARS-CoV-2 Spike protein was able to pseudotype both LV and VSV vectors, and determined that the combination of using a conditional VSV vector and a luciferase reporter gene had the advantage of allowing titers to be read at 16 hours post-transduction. Such Spike pseudotyped VSV vectors supported entry into a variety of mammalian cell types, including lung organoid systems, making them a useful system with which to study SARS-CoV-2 entry under standard laboratory conditions.

Optimization of pseudotyped vectors includes selection of an appropriate concentration method, such as centrifugation, PEG precipitation or ultrafiltration. For VSV pseudovectors, we found that concentration by TFF produced vector stocks with higher recovery rates and more consistent titers throughout a dilution series than those produced by ultracentrifugation. TFF also has the advantage of providing a partial purification due to the selective loss of potential contaminants below the cut-off value of the filter, and provides a larger processing capacity than ultracentrifugation. As a result, TFF is frequently used to facilitate large-scale vector production, including for clinical use (65, 66, 72). In our own experience, 3L of supernatant can be concentrated down to 50ml in 2 hours.

Since pseudovector titers can be impacted by incompatibilities between a viral fusion protein and the heterologous viral particle (45–50), an additional strategy to enhance vector titers has been to truncate the fusion protein’s cytoplasmic tail. We found that this approach increased the titers of Spike protein pseudovectors based on both LV and VSV, in agreement with previous reports (24, 59–63). Furthermore, as others have also noted (59, 62), the enhanced vector titers correlated with increased levels of Spike protein incorporation that were not simply the result of higher levels of cell surface expression following tail truncation, and tail truncation has also been reported to enhance the fusogenicity of Spike (59, 62). Together, this suggests that truncation of the cytoplasmic could also alter the conformation or function of the protein’s ectodomain, as has been reported for viral fusion proteins in HIV (57), measles virus (56), simian immunodeficiency virus (55) and gibbon ape leukemia virus (46), where truncation of the cytoplasmic tail impacted ectodomain conformation or functions such as receptor binding, or fusogenicity.

Our comparison of techniques to enhance vector titers also identified an area for caution; although cytoplasmic tail truncation enhanced pseudovector titers, they can also have unintended functional consequences. Specifically, we found that the impact of the D614G mutation on Spike protein incorporation and vector titer was obscured by the cytoplasmic tail truncation. A similar lack of effect of the D614G mutation on titer was also reported in another study using a 21 amino acid deletion of the Spike protein cytoplasmic tail in VSV pseudovectors (62). These findings suggest that tail-truncated Spike proteins should be used with caution for studies analyzing the impact of Spike mutations, or to test potential therapeutics targeting SARS-CoV-2 entry.

The mechanism for enhanced incorporation and/or titer by D614G is not entirely understood but structural analyses have suggested that it could impact Spike protein structure and stability, both within and between Spike monomers. For example, it has been suggested that a glycine at this location could strengthen the association between the S1/S2 subunits through an impact on the epistructure that decreases the intramolecular wrapping in the S1 subunit but promotes intermolecular wrapping between S1 and S2 (73). In an alternative model, the D614G mutation could alter the structure and/or stability of the Spike trimer by abrogating the hydrogen bond connecting D614 in the S1 subunit of one monomer with T859 in the S2 subunit of a neighboring monomer (34). These alterations were hypothesized to result in a greater tendency of the G614 monomers to form stable trimers which, in turn, could facilitate their incorporation into virions. As evidence, a mixture of equal amounts of D614 and G614 Spike variants expressed in vector-producing cells resulted in a higher level of G614 proteins in the incorporated Spike trimers (70).

Finally, we also used the VSV pseudovectors to evaluate the impact of the D614G mutation on infectivity of different cell types and sensitivity to antibody neutralization. Consistent with previous findings using pseudoviruses (32–34, 37, 43, 44) or SARS-CoV-2 virions (41, 68), we found that the G614 variant exhibited enhanced transduction of various cell lines when compared to the D614 variant, and that this correlated with increased Spike incorporation into the VSV particles (43, 70). As previously noted, these effects were significantly abrogated when tail truncated variants were used, consistent with an upper limit for the enhancement of Spike incorporation.

In contrast, the serum neutralization studies revealed no differences in sensitivity for either residue at position 614, and in either of the cytoplasmic tail configurations. This is in agreement with the majority of reports testing the impact of the D614G mutation with full-length Spike pseudovectors or SARS-CoV-2 virus against human convalescent serum (30, 34, 43, 68), serum from convalescent animals (41), or vaccinated human or animals (44, 68, 74).

In summary, we found that although cytoplasmic tail truncations enhance SARS-CoV-2 Spike protein incorporation into both LV and VSV vectors, and enhance the titers of unconcentrated vectors, they can also mask the phenotype of the D614G mutation. Pseudotyped vectors are increasingly being used to study newly emerging SARS-CoV-2 variants, where both full-length (42, 75) and truncated Spike proteins (25, 35, 39) have been used in studies investigating the impact of mutations on Spike protein properties such as ACE2 binding, transduction efficiency or sensitivity to neutralization. To better ensure the authenticity of the Spike protein functions being investigated in such vectors, we recommend using a full-length Spike protein, and combining vector production with TFF if higher titer vectors are required.

## METHODS

### Plasmids

Full-length (S) and 18 amino acid cytoplasmic tail truncated (SΔ18) Spike proteins for the Wuhan-Hu-1 isolate of SARS-CoV-2 (GenBank: MN908947.3) were provided by Dr. James Voss (The Scripps Research Institute) in a plasmid pcDNA3.3 backbone. D614G mutants were generated by site-directed mutagenesis. A VSV G protein expression plasmid was obtained from Addgene (Watertown, MA; Cat.# 8454).

### Cell lines

293T, HeLa, HeLa-ACE2, Vero, VeroE6 and Huh7.5 cells were maintained in Dulbecco’s modified Eagle medium (DMEM), and Calu-3 cells were maintained in Eagle’s Minimum Essential Medium (EMEM). All media were supplemented with 4 mM glutamine and 10% fetal bovine serum (FBS). HeLa-ACE2 cells were provided by Dr. James Voss, and were generated by transduction of HeLa cells with a lentiviral vector packaging a CMV-ACE2 expression cassette. The Huh7.5 cell line was provided by Dr. Jae Jung (Cleveland Clinic). All other cell lines were obtained from ATCC.

### VSV vector production, concentration and transduction

Replication-deficient VSVΔG vectors (76), containing expression cassettes for firefly luciferase or GFP in place of the VSV G protein, were provided by Dr. Jae Jung and Dr. Oscar Negrete (Sandia National Laboratories), respectively. To generate Spike pseudotyped VSV vectors, 4 × 10^6^ 293T cells were seeded in DMEM plus 10% FBS in a 10cm plate and transfected with 15 μg of Spike expression plasmid 24 hours later, using the calcium-phosphate transfection method (76). Media was replaced 16 hours later with 10 ml fresh media, and after a further 8 hours, 5 ml was removed and 2×10^8^ vector genomes of VSVΔG particles were added for one hour at 37 °C. Following this incubation, cells were washed three times with PBS and incubated for a further 24 hours before harvesting supernatants.

For larger scale production, quantities were adjusted to seed 3×10^7^ cells in 500cm^2^ plates, transfection with 124.5 μg of Spike expression plasmid and infection by 1.7×10^9^ vector genomes of VSVΔG particles per 500cm^2^ plate. To propagate VSVΔG particles, the same protocol was followed but replacing the Spike expression plasmid with the same quantity of a VSV G expression plasmid, and no PBS washes were performed after infection by VSVΔG.

Vector supernatants were harvested and filtered through 0.45 μm syringe filters, and either aliquoted or concentrated by ultracentrifugation using 20% (w/v) sucrose cushions for 2 hours at 25,000 rpm in an SW41 or SW28 rotor (Beckman, Indianapolis, IN). Alternatively, large-scale supernatant preparations were concentrated by tangential flow filtration (TFF) using a polyethersulfone membrane hollow fiber unit with 100 kDa molecular weight cut off and 155cm^2^ filtration surface (Spectrum Laboratories, Rancho Dominguez, CA) and a KR2i peristaltic pump (Spectrum Laboratories). To perform buffer exchange and prevent filter blockage, every 100 ml of vector supernatant was followed by 100 ml PBS. A 10- to 12-fold concentration from the original volume to approximately 8 ml final volume was achieved. All vectors were stored at −80°C in aliquots.

VSV-luciferase vector transductions were performed on tissue culture treated, 96-well half-area white plates (Corning, Corning, NY), seeded with various cells lines to achieve 50%-75% confluency at the time of transduction. Vectors were serially diluted and added to the culture to achieve final dilutions of 1:5, 1:15, 1:45, 1:135, and 1:405 and incubated at 37 °C for 16-24 hours. Transduction efficiency was quantified by measuring luciferase activity in cell lysates using Britelite Plus (Perkin Elmer, Richmond, California) and following the manufacturer’s protocol. To calculate the fold-change in transduction efficiency between D614 and G614 mutants, data from the 1:45 dilution points was used.

To titer VSV-GFP vectors, HeLa-ACE2 cells were seeding as 1×10^4^ cells per well in 96-well plates, and the following day, 50 μl of serially-diluted unconcentrated vector stocks were added. The final dilutions in the cultures were 1:2, 1:6, 1:18, and 1:27. Transduction efficiency was determined by GFP expression 16-24 hours after transduction using flow cytometry (Guava easyCyte, MilliporeSigma, Burlington, MA). and transducing units (TU) per ml calculated from the dilutions showing a linear relationship between the dilution factor and the number of GFP-positive cells.

### Lentiviral vector production, concentration and transduction

Lentiviral vectors were generated by transfection of 10 cm plates of 293T cells at 75% confluency with 2 μg of Spike expression plasmid, 10 μg of packaging plasmid pCMVdeltaR8.2 (Addgene Cat.# 12263) and 10 μg of a GFP-expressing vector genome plasmid FUGW (Addgene Cat.# 14883). Media was removed 16 hours later and replaced with 10 ml fresh DMEM plus 10% FBS. Supernatants were harvested 48 hours after transfection and filtered through 0.45 μm syringe filters, and either aliquoted or concentrated by ultracentrifugation using 20% (w/v) sucrose cushions for 2 hours at 25,000 rpm in an SW41 rotor (Beckman).

HeLa-ACE2 cells were transduced with Spike pseudotyped LV by seeding 1×10^4^ cells per well in 96-well plates and adding 50μl of unconcentrated vector stocks the next day. Transduction efficiency was determined by GFP expression 48 hours after transduction using flow cytometry, as described above, and reported as transducing units (TU) per ml.

### LV and VSV vector genome titration

RNA from 160 μl of LV or VSV vector stocks was extracted using Viral RNA mini kit (Qiagen, Hilden, Germany) and reverse transcribed into cDNA using SuperScript (Invitrogen, Carlsbad, CA), according to the manufacturer’s instructions. Genome copy number was determined by ddPCR for the WPRE sequences in the LV genome, or for Phosphoprotein (P) sequences in the VSV genome, using the QX200 Droplet Digital PCR system (Bio-Rad, Hercules, CA) and primer/probe set: WPRE-forward (CCTTTCCGGGACTTTCGCTTT), WPRE-reverse (GGCGGCGGTCACGAA), WPRE-probe (FAM- ACTCATCGCCGCCTGCCTTGCC-TAMRA), P-forward (GTCTTCAGCCTCTCACCATATC), P-reverse (AGCAGGATGGCCTCTTTATG), P-probe (FAM-TCGGAGGTGACGGACGAATGTCT-IOWA BLACK). Briefly, 6.25 μl of 1:10 and 1:100 diluted cDNA was mixed with forward and reverse primers (final concentration 900nM), probe (final concentration 250nM), 2x ddPCR supermix (Bio-Rad), and made up to 25 μl with water. Twenty microliters of each reaction mix was converted to droplets by the QX200 droplet generator, and droplet-partitioned samples were transferred to a 96-well plate and sealed. Thermal cycling was performed with the following conditions: 95 °C for 10 min., 40 cycles of 94 °C for 30 sec., 60 °C for 1 min., and 98 °C for 10 min. Plates were read on a QX200 reader (BioRad) and DNA copies quantified by detection of FAM positive droplets.

### Lung bud organoid differentiation and transduction

Lung bud organoids were generated from human pluripotent stem cells (hPSCs) and validated as previously described (77). hPSC differentiation into endoderm was performed in serum-free differentiation (SFD) medium of IMDM/Ham’s F-12 (3:1) (Life Technologies, Carlsbad, CA) supplemented with the following: 1 × N2 (Life Technologies), 0.5 × B27 (Life Technologies), 50 μg/ml ascorbic acid, 1 × Glutamax (Gibco), 0.4 μM monothioglycerol, 0.05% BSA, 10 μM Y27632, 0.5 ng/ml human BMP4 (R&D Systems), 2.5 ng/ml human FGF2 (R&D Systems, Minneapolis, MN), and 100 ng/ml human Activin (R&D Systems), in a 5% CO_2_/5% O_2_ atmosphere at 37 °C for 72-76 h. On day 4, endoderm yield was determined by the expressions of CXCR4 and c-KIT by flow cytometry. Cells used in all experiments had > 90% endoderm yield. For induction of anterior foregut endoderm, embryonic bodies were dissociated into single cells using 0.05% trypsin/0.02% EDTA and plated onto fibronectin-coated, six-well tissue culture plates (80,000–105,000 cells/cm^2^). Cells were incubated in SFD medium supplemented with 100ng/ml human Noggin (R&D Systems) and 10μM SB431542 for 24 hours followed by switching to SFD media supplemented with 10 μM SB431542 and 1 μM IWP2 (R&D Systems) for another 24 hours. At the end of anterior foregut endoderm induction, cells were maintained in SFD media supplemented with the following: 3 μM CHIR 99021 (CHIR, R&D Systems), 10 ng/ml human FGF10, 10 ng/ml human KGF, 10 ng/ml human BMP4 and 50nM all-trans retinoic acid for 48 hours, when three-dimensional cell clumps formed. Clumps were suspended by gently pipetting around the wells to form lung bud organoids, which were maintained in Ultra-Low Attachment multiple well plates (Corning) and fed every other day, and used for vector transduction after day 35.

To transduce lung bud organoids, 10 to 20 organoids were picked manually and transferred to 96-well U-bottom plates and transduced with 50 μl of GFP-expressing VSV vectors (1.7×10^4^ TU/ml). Transduction efficiency was examined by GFP expression 24 hours later by fluorescence microscopy.

### Western blot analysis of Spike protein incorporation

Vector supernatants were concentrated by ultracentrifugation (100-fold), electrophoresed on 4-12% Bis-Tris protein gels (Bio-Rad) and transferred to PVDF membranes using Trans-Blot Turbo Transfer System (Bio-Rad). Membranes were blocked with 5% milk in PBST buffer (PBS plus 0.1% of Tween®20). The S1 subunit of Spike was detected using SARS-CoV-2 (COVID-19) Spike S1 antibody at 1:1000 (Prosci, Cat.# 9083); the S2 subunit was detected using anti-SARS-CoV/SARS-CoV-2 (COVID-19) spike antibody clone [1A9] at 1:1000 (GeneTex, Cat.# GTX632604); HIV-1 p24 was detected using a polyclonal anti-HIV-1 SF2 p24 rabbit antiserum at 1:6000 (obtained through the NIH HIV Reagent Program, Division of AIDS, NIAID, NIH: ARP-4250, contributed by DAIDS/NIAID; produced by BioMolecular Technologies). VSV M protein was detected using anti-VSV M antibody clone [23H12] (KeraFast, Boston, MA, Cat.# EB0011) at 1:1000. HRP-conjugated goat anti-mouse and goat anti-rabbit antibodies were used as secondary antibodies (Santa Cruz Biotechnology, Dallas, TX). Blots were imaged by Amersham ECL Prime Western Blotting Detection Reagent (GE healthcare, Chicago, IL) and Chemidoc (Bio-rad). Densitometry was measured using ImageJ software (http://rsb.info.nih.gov/ij/).

### ACE2 cell surface expression by flow cytometry

HeLa-ACE2 cells were detached from culture flasks by 0.05% trypsin (Corning) and washed once with PBS. One million cells were re-suspend in 100 μl PBS and either immediately incubated with 0.25 μg anti-ACE2 antibody (R&D systems, Cat.# AF933) or first incubated at 37 °C for 6 hours with shaking to allow recovery of cell surface proteins after trypsinization. Alexa Fluor 647 conjugated donkey anti-goat antibody (1:200 dilution, Thermo Fisher Scientific, Waltham, MA, Cat.# A21447) was used as a secondary antibody and ACE2 expression was determined by flow cytometry (Guava easyCyte).

### Spike protein cell surface expression

VSV pseudovector-producing 293T cells were harvested to examine cell surface expression of the Spike protein. The S1 subunit was detected using SARS-CoV-2 (COVID-19) Spike S1 antibody at 1:100 (Prosci, Fort Collins, Colorado, Cat.# 9083) and the S2 subunit was detected using anti-SARS-CoV/SARS-CoV-2 (COVID-19) spike antibody clone [1A9] at 1:100 (GeneTex, Irvine, CA, Cat.# GTX632604). APC-conjugated goat anti-mouse and ducky anti-rabbit antibodies were used as secondary antibodies (1:100 dilution, Invitrogen), and expression was detected by flow cytometry (Guava easyCyte). The expression levels of different Spike proteins were reported as mean fluorescence intensity (MFI).

### Convalescent serum neutralization

Convalescent serum from COVID-19 patients or healthy donors (collected before April 30, 2020) was obtained from Children’s Hospital Los Angeles. Convalescent sera was confirmed to be positive for IgG class antibodies against SARS-CoV-2 Spike using anti-SARS-CoV-2 ELISA (IgG) (EUROIMMUN, Lübeck, Germany) (78).

A suitable dose of Spike pseudotyped VSV-Luc vectors was used in the neutralization assays to produce approximately 10^5^ relative light unit (RLU) of luciferase activity on HeLa-ACE2 cells in the absence of serum. Five ×10^3^ HeLa-ACE2 cells were seeded in tissue culture-treated, 96-well half-area white plates (Corning) to achieve 50%-75% confluency the following day. Convalescent or control sera were 3-fold serially diluted from 1:10 to 1:7290 and 50 μl incubated with the predetermined dose of the VSV-Luc vectors for 30 mins at 37°C, before addition of the mixture to HeLa-ACE2 cells. Cells were incubated at 37 °C overnight for 16-24 hours. Vector transduction efficiency was quantified by measuring luciferase activity as described above and neutralization (%) calculated by normalization to the values obtained on cells transduced without serum.

## ACKNOWLEDGMENTS

This work was supported by a grant to PMC from the W.M. Keck Foundation COVID-19 Research Fund to the Keck School of Medicine of USC. The authors would like to acknowledge the generosity of Drs. James Voss, Jae Jung and Oscar Negrete, who provided COVID-19 reagents, cell lines and VSVΔG particles.

